# Investigating TNS4 in the Colorectal Tumour Microenvironment Using 3D Spheroid Models of Invasion

**DOI:** 10.1101/2020.01.03.893958

**Authors:** Teresa P. Raposo, Susanti Susanti, Mohammad Ilyas

**Author notes:** Correspondence to: Teresa Pereira Raposo,: West block D floor, Cancer Biology, Queen’s Medical Centre, Derby Road, NG7 2UH Nottingham, United Kingdom.

## Abstract

TNS4 (Tensin 4 or Cten) is a putative oncogene in colorectal cancer (CRC) with a role in regulating cell adhesion, motility, invasion and epithelial to mesenchymal transition (EMT). Our objective was to study TNS4 role in CRC using more realistic models of the tumour microenvironment.

CRC cells expressing TdTomato protein and shTNS4/shLUC hairpin oligos were grown in 3D spheroids with and without cancer-associated fibroblasts (CAFs). Adhesiveness to collagen I and CAFs was assessed in 2D and cell proliferation, volume and invasion were assessed in 3D conditions. The role of TNS4 knockdown in Gefitinib chemosensitivity and EGFR and Ras protein levels were also tested.

In general, TNS4 knockdown increased cell proliferation in cell lines producing compact spheroids. The addition of CAFs in spheroids supported CRC cells proliferation, whereas CAFs themselves did not proliferate, but increased ECM degradation. TNS4 knockdown reduced adhesiveness and 3D invasion and disrupted EGFR signalling which resulted in increased sensitivity to Gefitinib.

In conclusion, in a 3D spheroid model, TNS4 inhibits cell proliferation and promotes cell invasion into the ECM, possibly by adhesion to the ECM and stromal cells. TNS4 knockdown enhances sensitivity to the EGFR inhibitor Gefitinib and may be helpful for KRAS mutant CRC patients.

## Introduction

TNS4 (tensin4) or Cten (C-terminal tensin-like) is a tensin localized to focal adhesion sites where it has a role in the dynamics of cell movement and adhesion by linking integrins and scaffold proteins to the actin cytoskeleton and the extracellular matrix (ECM) [1, 2].

During the past decade, TNS4 has been suggested as a putative oncogene due to its overexpression in a vast range of cancer types: breast, colorectal, hepatocellular carcinoma, melanoma, oesophagogastric adenocarcinoma, gastric, pancreatic, thymoma and lung cancer [3–11].

Even though TNS4 has been identified as an oncogene in a number of cancers, the mechanisms regulating its role in motility, invasion and metastasis are still under investigation.

In CRC cell lines, TNS4 has been shown to induce EMT by repressing E-cadherin and to significantly increase migration and invasion [3]. More recently, a role for TNS4 in selectively mediating TGF-ß-induced EMT in colorectal cancer cell lines has been reported. In the absence of TNS4, the effects of TGF-ß stimulation on inducing motility and invasion were abrogated, but TGF-ß-induced proliferation was not affected [12]. Furthermore, TNS4-mediated Src stabilization was found to be the responsible for EMT induction in CRC cell lines [13] Nude mice receiving a splenic injection of TNS4 overexpressing HCT116 cells have produced larger splenic tumours and hepatic metastatic nodules compared to the control group [5]. However, thus far, studies demonstrating the role of TNS4 in syngeneic or immunocompetent animal models of colorectal cancer invasion and metastasis *in vivo* are still lacking.

The use of 3D *in vitro* models may fill in this gap by providing an enriched tumour microenvironment with improved cell interactions and ECM which better mimic the *in vivo* context and which may provide a better measure of drug response [14–18]. Addition of stromal cell types such as cancer-associated fibroblasts (CAFs) to 3D models substantially supports growth of cancer cell lines and may also promote invasion and metastasis by guiding cancer cells at the invasive edge [19]. Interactions between tumour and stroma are difficult to study *in vivo* due to limitations in tissue accessibility, *in vivo* monitoring and selection of specific intercellular interactions [20].

In this study we aim firstly to explore the role of TNS4 in heterotypic interactions between cancer cells and CAFs within the tumour spheroid and in an *in vitro* tumour microenvironment, and secondly to validate the role of TNS4 in colorectal cancer proliferation and invasion by using a 3D spheroid model of the tumour microenvironment. To this end we have used stable lentiviral TNS4 and Luciferase control knockdowns of colorectal cancer cell lines expressing TdTomato fluorescent protein, alone or with cancer-associated fibroblasts in mixed spheroids. Expression of fluorescent TdTomato was used to measure the growth of colorectal cancer cells within the spheroid and GFP-tagged CAFs allowed imaging of CAFs invasion in the spheroids.

## Materials and methods

### Cell culture

CAFs were isolated from human colon carcinoma resection specimens following 2.4U/ml dispase + 100U/ml collagenase type II (both from Invitrogen); disaggregation of tumour fragments and digestion for 2h t 37°C. Cells were cultured in DMEM (Gibco) supplemented with 20% heat-inactivated FBS (Gibco), 2mM L-glutamine, 0.1% hydrocortisone (w/v), 0.75% bovine insulin (w/v).

All cell lines used in this study were obtained from ATCC and used for experiments up to passage 30. SW620^ΔCten^ is derived from SW620 submitted to TNS4 deletion by CRISPR-Cas9 [21].

HEK293T, DLD1, HCT116, SW480, SW620, SW620^ΔCten^ were maintained in 2D cell culture in DMEM (Gibco) supplemented with 10% heat-inactivated FBS (Gibco), 2mM L-glutamine and incubated at 37°C, 5% CO_2.._ Cells were regularly passaged by dissociation with Trypsin-EDTA 0.25% (Sigma Aldrich) every 2-3 days.

HT29 was maintained in McCoys 5A medium supplemented with 10% FBS and 2mM L-glutamine. LS1034 was maintained in RPMI 1640 supplemented with 10% FBS and 2mM L-glutamine. Gefitinib (Sigma Aldrich, Y001813) was diluted to the concentration of 100mM in DMSO and subsequently dissolved in cell culture media to concentrations of 50 and 25 µM. Cell viability was measured in six replicates, by using a Fluorostar Omega plate fluorescence plate reader (540 nm excitation, 590 nm emission) after incubation with 10% Presto Blue (Life Technologies) for 1h at 37C, 5%CO2.

### Lentivirus production and transduction

The lentiviral plasmid components were acquired from Addgene: Transfer plasmids: Rix-PGK-Tom-W (#25813), FUGW (#14883) and PLKO.1 TRC (# 10878), Packaging: psPAX2 (#12260) and Envelope: pMD2.G (#12259).

The short hairpin RNA (shRNA) for TNS4 and LUC (Luciferase) knockdown were selected from the RNAi consortium shRNA library (https://www.broadinstitute.org/rnai-consortium/rnai-consortium-shrna-library).

The shRNA oligos (Supplementary Table 1) were annealed at 95°C for 4 min in a PCR thermal cycler and slowly cooled down overnight. The annealed oligos were phosphorylated, ligated into pLKO.1-TRC, transformed in chemocompetent E.coli (New England Biolabs), isolated by miniprep and double-digested with restriction enzymes AgeI and EcoRI (both New England Biolabs). Correctly inserted oligos were confirmed by restriction enzyme double-digestion with EcoRI and NcoI of the resulting vectors and sequencing.

Viral particles were packaged in HEK293T cells seeded at a density of 1.5×10^6^/10cm petri dishes in a proportion of 24μL Fugene 6 (Promega, E2691) in 136μL serum-reduced OPTI-MEM (Gibco, Invitrogen) to 2μg pLKO.1 shRNA plasmid: 1500ng psPAX2 packaging plasmid (Addgene #12260): 500ng pMD2.G envelope plasmid (Addgene #12259) to 40μL serum-reduced OPTI-MEM. 72h after transfection, shRNA lentiviruses were concentrated by ultracentrifugation at 10,000xg, 4°C, 4h in a 10% sucrose buffer [22]. Lentivirus titers were examined by Lenti-go stix quick test (Takara Bio, 631280) according to the manufacturers’ instructions, using the freely mobile app (Lenti-X GoStix) to determine titer concentrations.

Colorectal cancer cell lines were firstly transduced with PLKO.1 shTNS4/ shLUC lentivirus and selected with Puromycin (Sigma Aldrich, p8833) at a dose that killed 100% of non-transduced control cells over a 7-day period, followed by transduction with TdTomato lentivirus at excessive multiplicity of infection (MOI) to ensure complete transduction efficiency. Cancer-associated fibroblasts were transduced with FUGW lentivirus to induce EGFP expression also at excessive MOI. Maintenance of complete transduction efficiency was routinely inspected by observation of TdTomato and EGFP expression across several passages, using a wide-field Nikon TiE fluorescence microscope.

### Protein extraction and Western blot

Cells were washed with ice-cold PBS and incubated on ice for 10min with RIPA buffer (Invitrogen) with protease and phosphatase inhibitor (Pierce, ThermoFisher) diluted to 1x. Resulting cell lysates were centrifuged at 16000g, at 4°C for 30 min and the supernatant transferred to a new eppendorf tube. A BCA assay was performed to determine protein quantification of the supernatant.

For Western blotting, 40µg of protein mixed with 25%(v/v)NuPAGE® LDS Sample buffer(4x) and 10%(v/v)reducing sample buffer (10x) were denatured at 95°C for 5 min, incubated in ice for 5 min and then loaded into to a pre-cast gel (NuPAGE® 4-12% Bis-Tris Protein Gels, Invitrogen). Gel electrophoresis was performed for 90 min at 125V. The resulting gel was immunoblotted onto a nitrocellulose membrane (Amersham, GE Healthcare Life Sciences) at 25V for 30 min, using the Trans-Blot Transfer system (BIO-RAD®).

After confirming transfer by Ponceau Red staining, the membrane was blocked with 5% skimmed milk (Sigma-Aldrich) in tris-buffered saline (TBS) for 1 hour with gentle agitation at room temperature. The membrane was then incubated overnight with primary antibodies at 4°C with gentle agitation (dilutions and suppliers listed on supplementary table 2) and then washed 3x 5min with TBST. The hybridised membrane was incubated with the respective IRDYE-conjugated secondary antibody (Donkey anti-mouse or anti-rabbit 800CW) for 1 hour at room temperature, using the blocking solution as the diluent. The membrane was further washed 3x 5min in TBST and infra-red fluorescence detected with the Li-Cor Oddissey® Blot scanner. Densitometry of the western blot bands was performed with Image studio lite software (LI-COR). Original imaged blots are shown in the negative image of the captured fluorescence light for better contrast without incurring in any modifications that could biase the perceived band intensity.

### 3D Spheroid formation, volumetry and fluorescence readings

Cell suspensions of the TdTomato expressing colorectal cell lines, their shTNS4 and shLUC expressing variants, were seeded in 12-replicates in Ultra-low attachment 96-well plates (CLS4515, Corning) at 2.0×10^4^cells/mL in a final volume of 100µL alone or in a 1:1 mix with CAFs, in Fluorobrite DMEM supplemented with 10% FBS and 4mM L-glutamine. The plates were centrifuged at 200g for 5 min, at acceleration 7 and deceleration 7 in a Megafuge 16 centrifuge (ThermoScientific). Spheroid or aggregate formation was confirmed in 24 to 48h and daily measurements of fluorescence were taken in a Fluostar Omega plate reader (BMG labtech) set to perform bottom readings, at 544nm excitation and 590 nm emission wavelength, gain 2500. Fluorescence readings were normalised to day 1 to account for initial differences in seeded cell numbers. Brightfield images of spheroids were taken daily for volumetry quantification using a widefield inverted microscope (Nikon TiE). Volumetry analysis was performed using Image J Wand tracing tool in 8-connected mode to determine the area of each spheroid. Mean spheroid radius was determined by assuming an approximation to the area of the circle (*A* = *πr*^2^) and from there inferred the volume of the spheroid 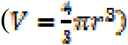.

### Adhesion assays

Adhesion assays to collagen type I and CAFs were performed to assess adhesiveness of shTNS4 cells versus its shLUC control. 96-well flat bottom plates (Costar) were coated with 100uL of 0.3mg/mL of rat tail collagen type I (Corning) diluted in PBS, 0.02M NaHO. Plates were incubated overnight at 37°C. For adhesion to CAFs, 96 well plates were seeded with 1000 CAFs/well and incubated overnight at 37°C. CRC cell lines were added to coated or CAF seeded plates at a density of 2.0×10^4^cells/well, and incubated for 2h. Half of the wells containing shTNS4 and shLUC cells were washed 3x with PBS and then incubated with complete fluorobrite DMEM for 15 min. Fluorescence values at 544-590nm excitation-emission were read in a Fluostar Omega plate reader. After subtracting background fluorescence, a ratio of washed to non-washed was used to determine cell adhesion.

### 3D spheroid invasion assay

At day 3 of spheroid formation, spheroids were transfered to a chilled flat-bottomed 96-well plate coated with 50uL/well of Cultrex BME RGF (Trevigen) at 6mg/mL and overlaid with another 50uL/well of Cultrex and 100uL of complete Fluorobrite DMEM supplemented with 80ng/mL human recombinant EGF as a chemoattractant. For extracellular matrix degradation analysis, DQ BSA-Green was added to Cultrex at 30ug/mL as previously described [23].

Spheroids were imaged at 96h of invasion by confocal microscopy, performing Z-stacks with 3um thick slices using 488nm and 568nm lasers. The volume of each fluorescent component was calculated on Image J given by multiplying the thresholded area (obtained by the Otsu method on the Image J threshold menu options) of each slice by its thickness, using a macro plugin freely available from http://www.optinav.info/MeasureStack.htm. A ratio of the measured DQ BSA green per TdTomato fluorescent volume was used as an estimation of ECM degradation normalised to the volume of cancer cells.

### Statistical analysis

Analysis of proliferation and volumetry assays was performed on GraphPad Prism version 7 by 2-way ANOVA with multiple mean comparisons by Tukey test. Comparisons between two groups of non-parametric values were performed by Mann-Whitney test. Statistical significance was considered for values of P<0.05.

## Results

### Stable TNS4 knockdown CRC cell lines

A stable knockdown of TNS4 was achieved by lentiviral transduction of colorectal cell lines (Figure 1) following which, transduction with TdTomato lentivirus allowed for tracking the growth of the CRC cell population within the spheroids by using a conventional fluorescent plate reader. I

**Figure 1.**
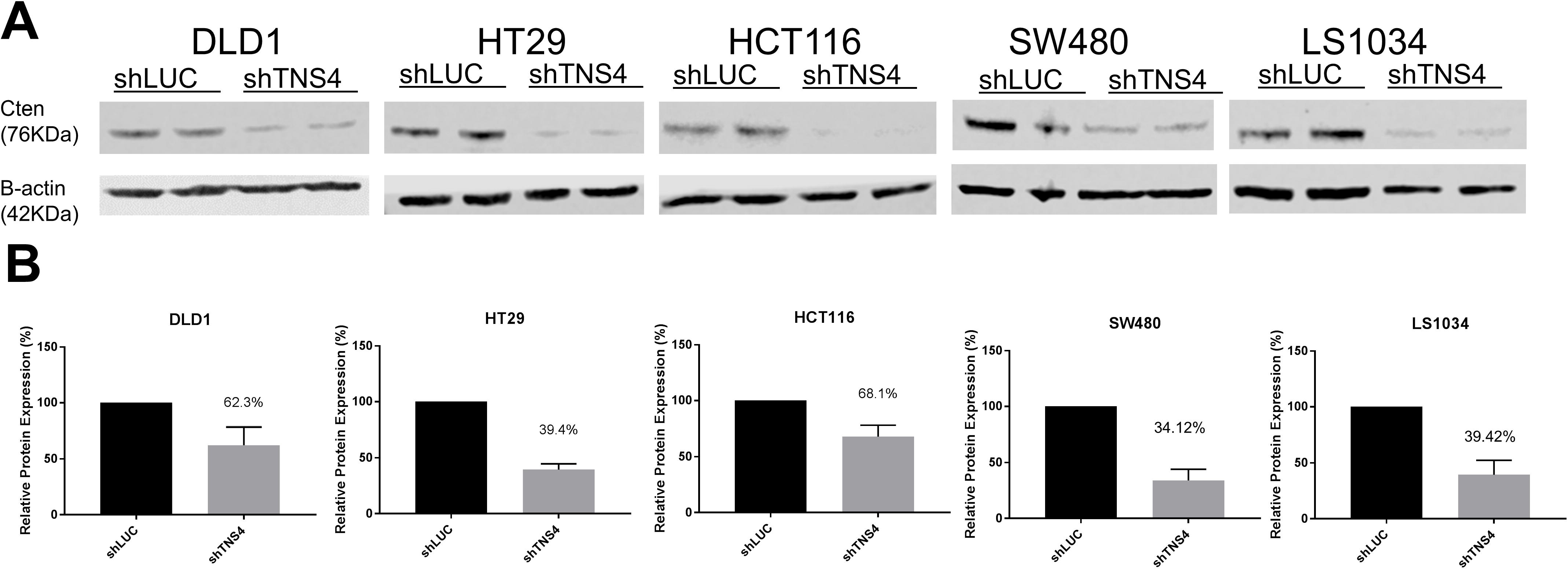
A: Western blot and respective densitometry showing a decrease in TNS4 expression achieved for shTNS4 stable knocdowns in DLD1, HT29, HCT116, SW480 and LS1034 cell lines relatively to their shLUC (luciferase control). Two independent protein lysates were tested. B: Densitometry ratios to the loading control B-actin, normalised to the shLUC levels (100%).

**Figure 2.**
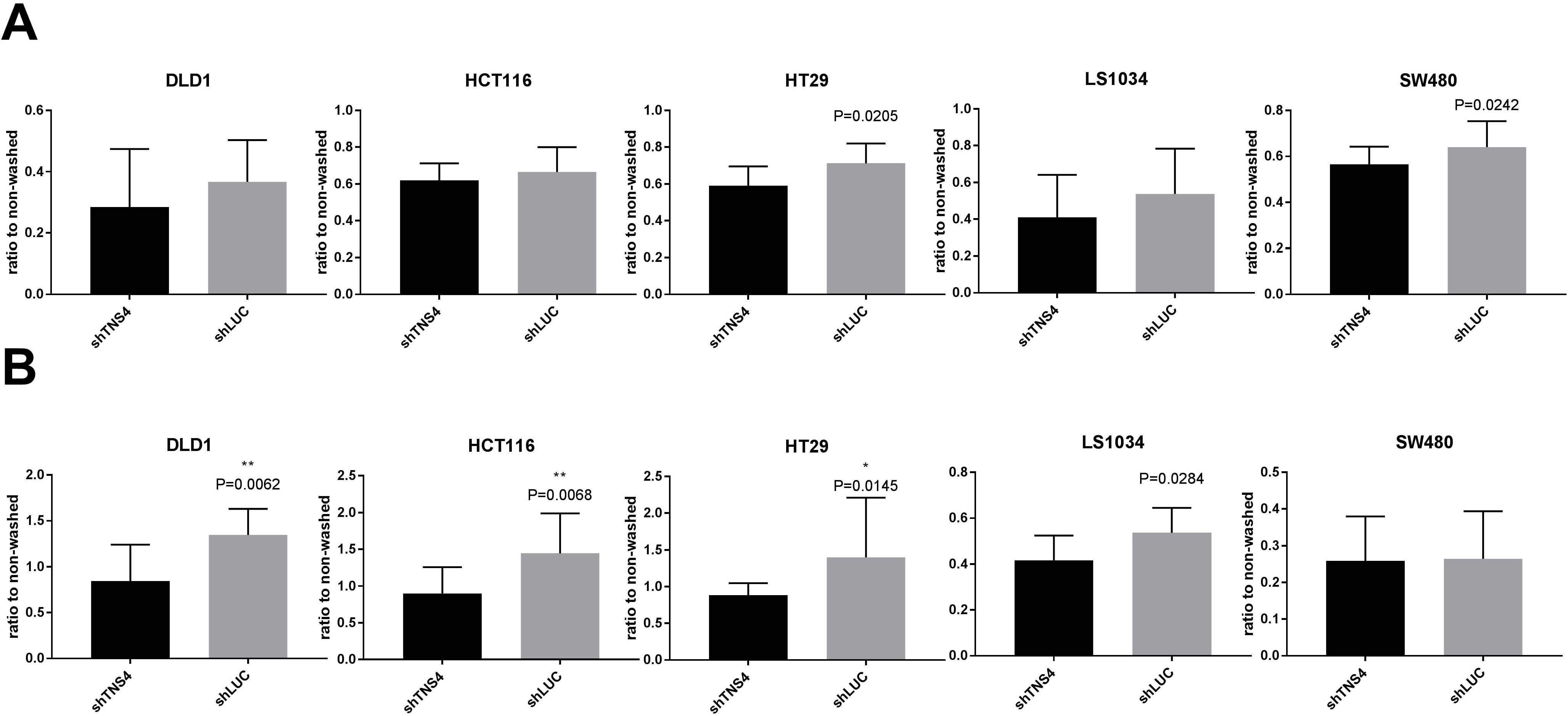
Adhesion assay of shTNS4 and shLUC CRC cells to collagen type I (A) and CAFs feeder layer (B). Mann-Whitney statistical test was performed to evaluate differences between shTNS4 and shLUC CRC cells. N=12 replicates and results shown are representative of 2 independent experiments. Statistically significant P-values are presented on top of the respective shTNS4 bars, in cases where statistically significant differences were not present, irrelevant P-values are omitted.

The spheroid formation method used revealed different morphologies between CRC cell lines. Whilst DLD1, HT29 and HCT116 formed compact spheroids of round uniform shape, SW480, and SW620 produced aggregates and LS1034 produced a globous irregular shape. (Suplementary Figure 1).

### TNS4 knockdown reduces adhesiveness

In the attempt to explain the observed differences in spheroid size and growth we have assessed the ability of shTNS4 and shLUC CRC cells to adhere to ECM collagen type I, which would normally be secreted by CAFs, or CAFs themselves, by performing adapted adhesion assays in collagen type I coated plates or using CAFs as an adhesion substrate. Whereas a tendency for reduced adhesion to collagen type I could be observed for all shTNS4 CRC cell lines, the difference to shLUC controls was only significant in HT29 (P=0.0205) and SW480 (P=0.0245) (Figure 4 A). By repeating the adhesion assay over a layer of CAFs, statistically significant reduced attachment of shTNS4 vs shLUC was noted for DLD1 (P=0.0062), HCT116 (P=0.0068), HT29 (P=0.0145) and LS1034 (P=0.0284).

The reduced adhesion to CAFs upon TNS4 knockdown could have an effect in the expansion and paracrine growth signalling depending on cell-to-cell contact between CAFs and CRC cells, leading to increased fluorescence signal in shTNS4 vs shLUC +CAFs spheroids. Conversely, a reduced spheroid volume seen in shTNS4+CAFs compared to shLUC+CAFs could be caused by increased compactness of the CRC cells, unsupported by a CAFs scaffold.

### TNS4 knockdown in 3D spheroids affects proliferative rates

To assess the growth of the CRC cell population within the spheroid, TdTomato fluorescence was measured daily over 7 days.

The SW620^ΔCten^ variant, in which TNS4 was deleted by using a CRSIPR-Cas9 system, consistently showed increased spheroid growth when compared to its parental SW620 cell line (P<0.0001) (Figure 1A, Table 1A). For both SW480 and LS1034 cell lines, the shTNS4 variant spheroids showed significantly reduced growth when compared to shLUC control spheroids (Figure 1 B,C). In DLD1, HCT116 and HT29, shTNS4 variants produced spheroids with slightly increased proliferation, but which was only significant for HCT116 and HT29 (P<0.0001) (Figure 1D, E, F and Table 1D,E,F).

**Table 1.** Statistical analysis of TdTomato mean fluorescence readings at day 7 for spheroids composed of CRC cell lines with and without CAFs. For each condition N=12 replicates and results are representative of 2 experiments. 2-way ANOVA statistical test with multiple comparisons by Tukey’s test was performed, a P-value <0.05 was considered statistically significant.

| **TdTomato relative fluorescence multiple comparisons at Day7** |  |  |
| --- | --- | --- |
| **A)** | **Mean Diff.** | **Adjusted P Value** |
| SW620 vs. SW620+CAF<sub>s</sub>(1:1) | -2.085 | <0.0001 |
| SW620 vs. SW620<sup>DCten</sup> | -0.8601 | <0.0001 |
| SW620<sup>DCten</sup> vs. SW620<sup>DCten</sup>+CAF<sub>s</sub> | -1.022 | <0.0001 |
| SW620+CAF<sub>s</sub> vs. SW620<sup>DCten</sup>+CAF<sub>s</sub> | 0.2036 | 0.4229 |
| **B)** |  |  |
| SW480 shTNS4 vs. SW480 shTNS4+CAF<sub>s</sub> | -0.1476 | <0.0001 |
| SW480 shTNS4 vs. SW480 shLUC | -0.209 | <0.0001 |
| SW480 shTNS4+CAF<sub>s</sub> vs. SW480 shLUC+CAF<sub>s</sub> | -0.08405 | 0.0197 |
| SW480 shLUC vs. SW480 shLUC+CAF<sub>s</sub> | -0.02262 | 0.8611 |
| **C)** |  |  |
| LS1034 shTNS4 vs. LS1034shTNS4+CAF<sub>s</sub> | 0.09316 | 0.0526 |
| LS1034 shTNS4 vs. LS1034 shLUC | -0.1275 | 0.0029 |
| LS1034shTNS4+CAF<sub>s</sub> vs. LS1034 shLUC+CAF<sub>s</sub> | -0.05505 | 0.4299 |
| LS1034 shLUC vs. LS1034 shLUC+CAF<sub>s</sub> | 0.1657 | <0.0001 |
| **D)** |  |  |
| DLD1 shTNS4 vs. DLD1 shTNS4+CAF<sub>s</sub> | -0.322 | <0.0001 |
| DLD1 shTNS4 vs. DLD1 shLUC | 0.03376 | 0.4726 |
| DLD1 shTNS4+CAF<sub>s</sub> vs. DLD1 shLUC +CAF<sub>s</sub> | 0.1275 | <0.0001 |
| DLD1 shLUC vs. DLD1 shLUC +CAF<sub>s</sub> | -0.2283 | <0.0001 |
| **E)** |  |  |
| HCT116 shTNS4 vs. HCT116 shTNS4+CAF<sub>s</sub> | -0.04509 | 0.8057 |
| HCT116 shTNS4 vs. HCT116 shLUC | 0.4322 | <0.0001 |
| HCT116 shTNS4+CAF<sub>s</sub> vs. HCT116 shLUC+CAF<sub>s</sub> | 0.3812 | <0.0001 |
| HCT116 shLUC vs. HCT116 shLUC+CAFs | -0.0961 | 0.2243 |
| **F)** |  |  |
| HT29 shTNS4 vs. HT29 shTNS4+CAFs | 0.0251 | 0.9859 |
| HT29 shTNS4 vs. HT29 shLUC | 0.8126 | <0.0001 |
| HT29 shTNS4+CAFs vs. HT29 shLUC+CAFs | 0.2259 | 0.0176 |
| HT29 shLUC vs. HT29 shLUC+CAFs | -0.5616 | <0.0001 |

**Table 2.**
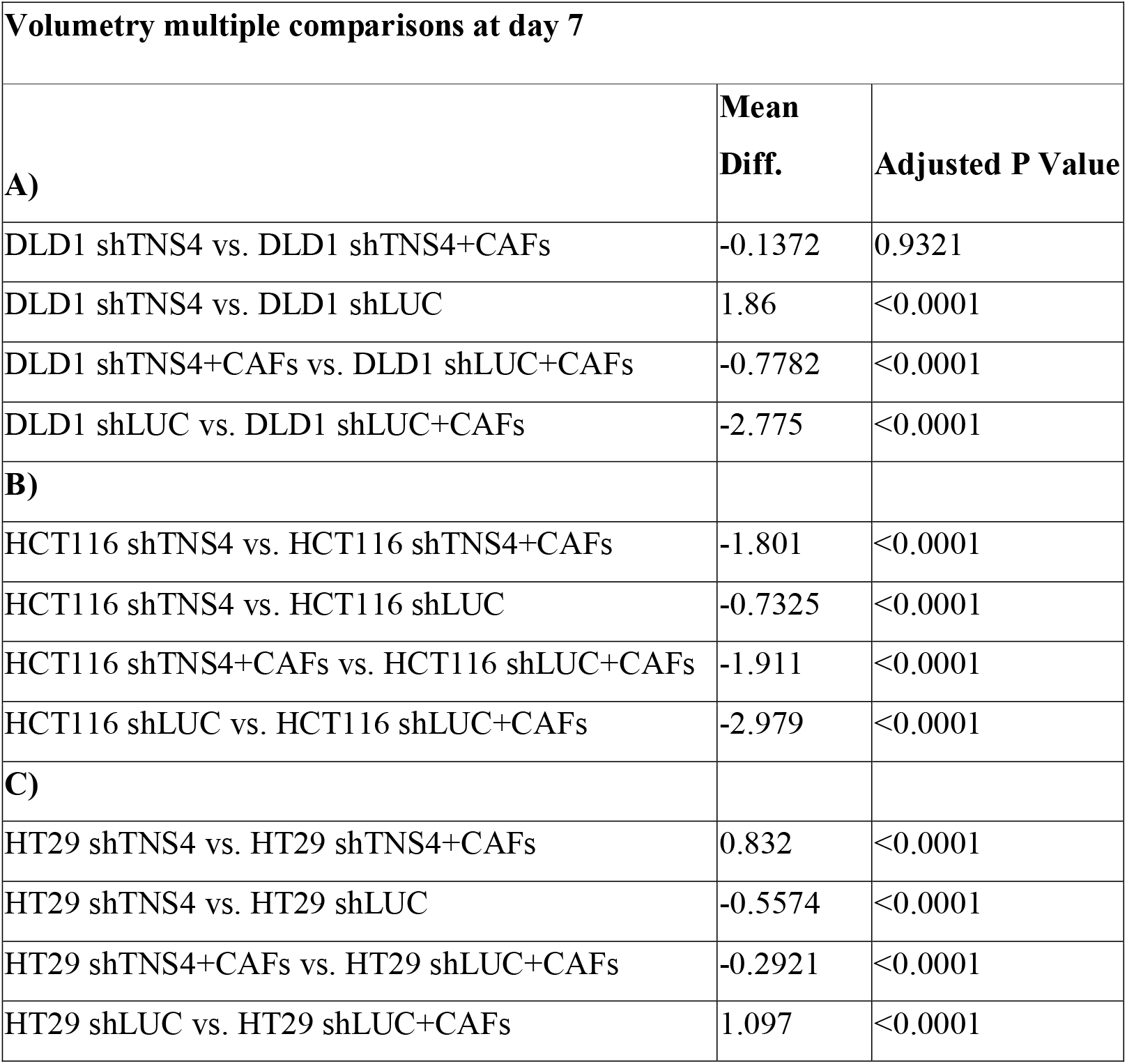
Statistical analysis of volume measurements at day 7 for spheroids composed of CRC cell lines with and without CAFs. For each condition N=12 replicates and results are representative of 2 experiments. 2-way ANOVA statistical test with multiple comparisons by Tukey’s test was performed, a P-value <0.05 was considered statistically significant.

| **Volumetry multiple comparisons at day 7** |  |  |
| --- | --- | --- |
|  | **Mean**<br>**Diff.** | **Adjusted P Value** |
| **A)** |  |  |
| DLD1 shTNS4 vs. DLD1 shTNS4+CAF<sub>s</sub> | -0.1372 | 0.9321 |
| DLD1 shTNS4 vs. DLD1 shLUC | 1.86 | <0.0001 |
| DLD1 shTNS4+CAF<sub>s</sub> vs. DLD1 shLUC+CAF<sub>s</sub> | -0.7782 | <0.0001 |
| DLD1 shLUC vs. DLD1 shLUC+CAF<sub>s</sub> | -2.775 | <0.0001 |
| **B)** |  |  |
| HCT116 shTNS4 vs. HCT116 shTNS4+CAF<sub>s</sub> | -1.801 | <0.0001 |
| HCT116 shTNS4 vs. HCT116 shLUC | -0.7325 | <0.0001 |
| HCT116 shTNS4+CAF<sub>s</sub> vs. HCT116 shLUC+CAF<sub>s</sub> | -1.911 | <0.0001 |
| HCT116 shLUC vs. HCT116 shLUC+CAF<sub>s</sub> | -2.979 | <0.0001 |
| **C)** |  |  |
| HT29 shTNS4 vs. HT29 shTNS4+CAF<sub>s</sub> | 0.832 | <0.0001 |
| HT29 shTNS4 vs. HT29 shLUC | -0.5574 | <0.0001 |
| HT29 shTNS4+CAF<sub>s</sub> vs. HT29 shLUC+CAF<sub>s</sub> | -0.2921 | <0.0001 |
| HT29 shLUC vs. HT29 shLUC+CAF<sub>s</sub> | 1.097 | <0.0001 |

Regarding the influence of CAFs in spheroid growth, a statistically significant increase in the growth of CRC cells in the spheroids was observed in SW620, SW480 and DLD1 when comparing shTNS4 variants with or without added CAFs in a 1:1 proportion (Table 1). Interestingly, in HT29 and LS1034, the promoting effect of CAFs was not seen in shTNS4 spheroids, but only in the presence of normal TNS4 expression in the shLUC spheroids (Table 1).

When comparing shTNS4 and shLUC co-cultured with CAFs, a significant growth boosting effect can be perceived for DLD1, HCT116 and HT29, in which spheroids composed of shTNS4 CRC cells with CAFs grow more than their control spheroids shLUC with CAFs. In CRC cell lines producing loose spheroid aggregates (SW620, SW480 and LS1034) this effect is not observed (Figure 3 and Table 1).

**Figure 3.**
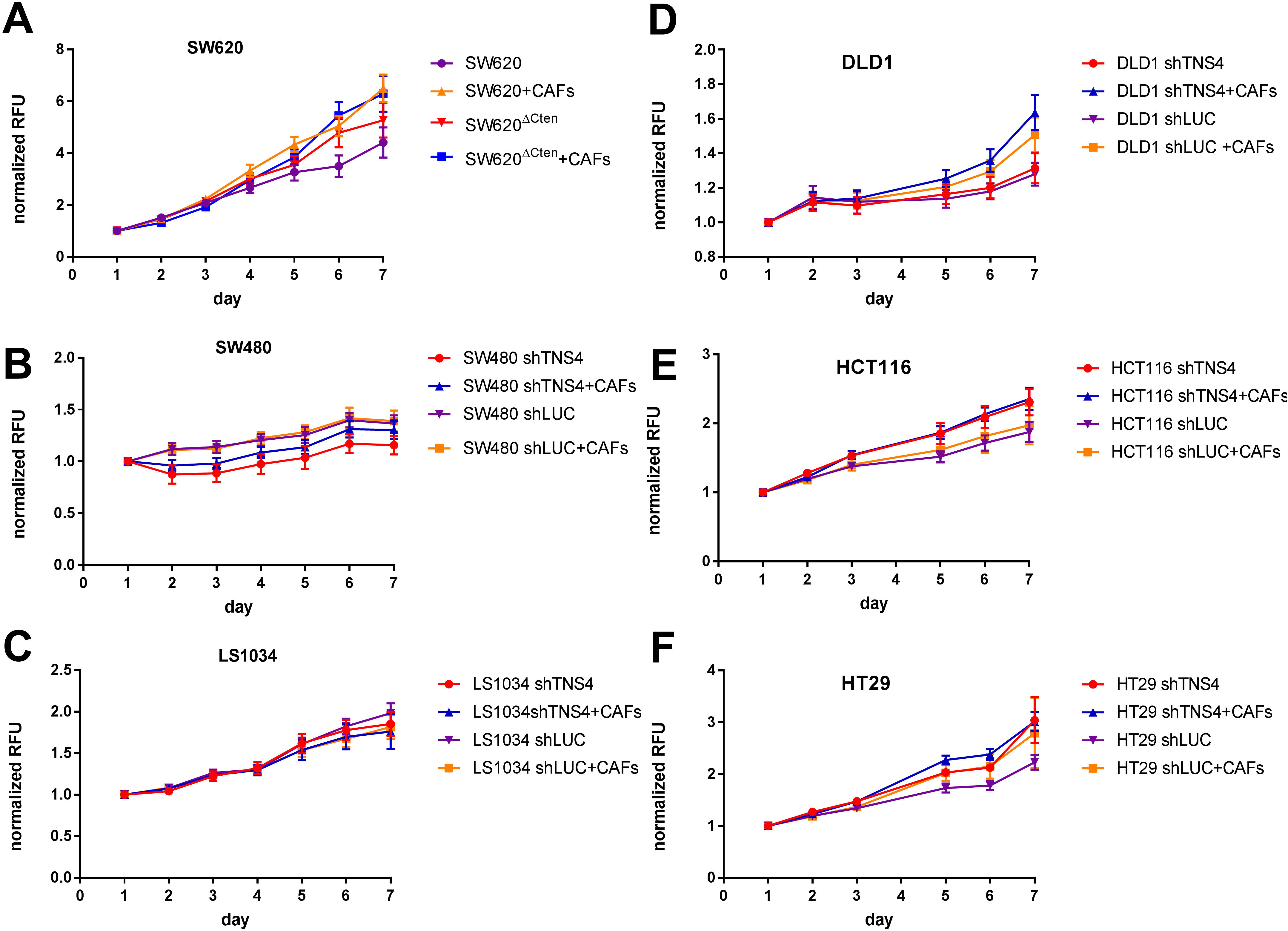
Mean+/− St dev of relative TdTomato fluorescence read-outs (540em-590ex nm) for spheroids composed of colorectal cell lines stably knockdown for TNS4 (shTNS4), luciferase control (shLUC) with and without addition of cancer-associated fibroblasts (CAFs). Graphs are representative of 12 replicates per condition and 2 independent experiments were carried out (the respective statistical analysis for this figure is shown on table 1).

**Figure 4.**
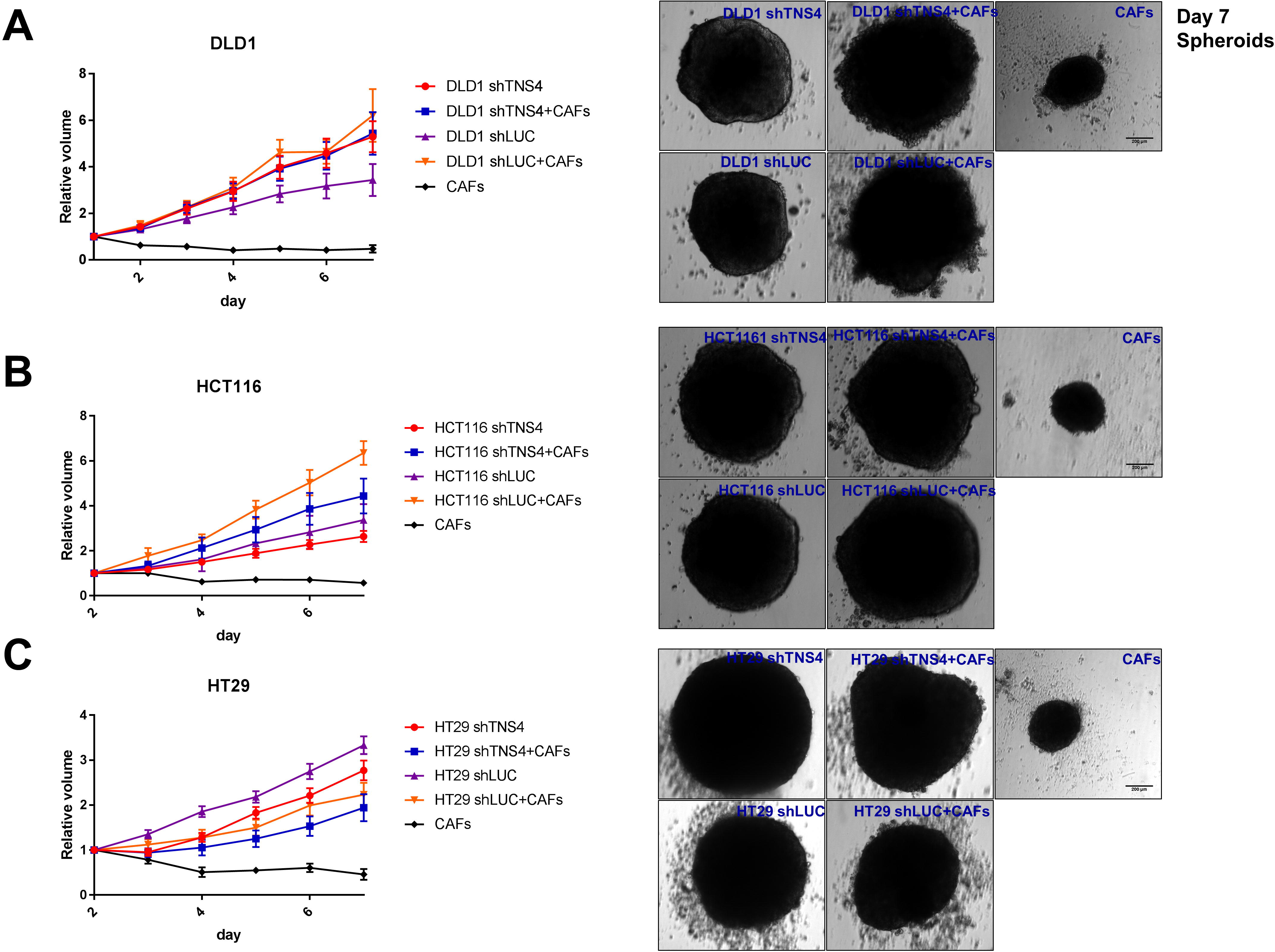
Volumetric measurements of spheroid growth in DLD1 (A), HCT116 (B) and HT29 (C) CRC cell lines and representative images of spheroids composed of shTNS4/shLUC variants with or without added CAFs at day 7.(The respective statistical analysis for this figure is shown on table 2).

Regarding differences observed in terms of whole spheroid volume, these could only be measured in those cell lines forming round compact spheroids (DLD1, HCT116 and HT29), in which the estimation made by approximation to the volume of a sphere is valid (Supplementary Figure 1 illustrates the spheroid morphology in the different cell lines used in this study). To normalise volumetric data, measurements taken on the second day were chosen for HCT116 and HT29, as spheroids of these cell lines only acquire a compact round shape later than DLD1. In HCT116 shTNS4 and shLUC variants as well as in DLD1 shLUC, addition of CAFs promoted spheroid growth in volume, only the difference between DLD1 shTNS4 and DLD1 shTNS4+CAFs was not statistically significant (Figure 3A, Table 2). However, in HT29, addition of CAFs seemed to reduce whole spheroid volume, possibly by contraction or increased compactness of the spheroid around CAFs (Figure 3C). When grown in low adherence conditions, CAFs form small spheroids and do not expand their volume, apparently in a quiescent state (Figure 3).

### TNS4 knockdown reduced 3D invasion and response to CAFs-driven invasion

By performing 3D invasion assays measuring specific ECM degradation without quenching of DQ BSA green we were able to isolate matrix invasion from changes in spheroid size due to a variation in cell proliferation.

Comparing CRC spheroids, there was a significant decrease in 3D invasion in shTNS4 versus shLUC spheroids in both DLD1 (P=0.0434) and HCT116 (P=0.0281), but not in HT29 (Figure 5A). In general, addition of CAFs to form mixed population spheroids increased ECM degradation and invasion, although this was not always a statistically significant effect. An exception to CAFs’ promotion of ECM degradation and invasion was observed with HT29 shTNS4 cells (Figure 5A and D), whereas a significant effect was observed for HT29 shLUC vs HT29 shLUC+CAFs. Interestingly, HT29 was the only CRC cell line where matrix degradation and invasion in shLUC+CAFs was superior to that of shTNS4+CAFs cells, suggesting a particular effect of TNS4 expression in the interaction with CAFs.

**Figure 5.**
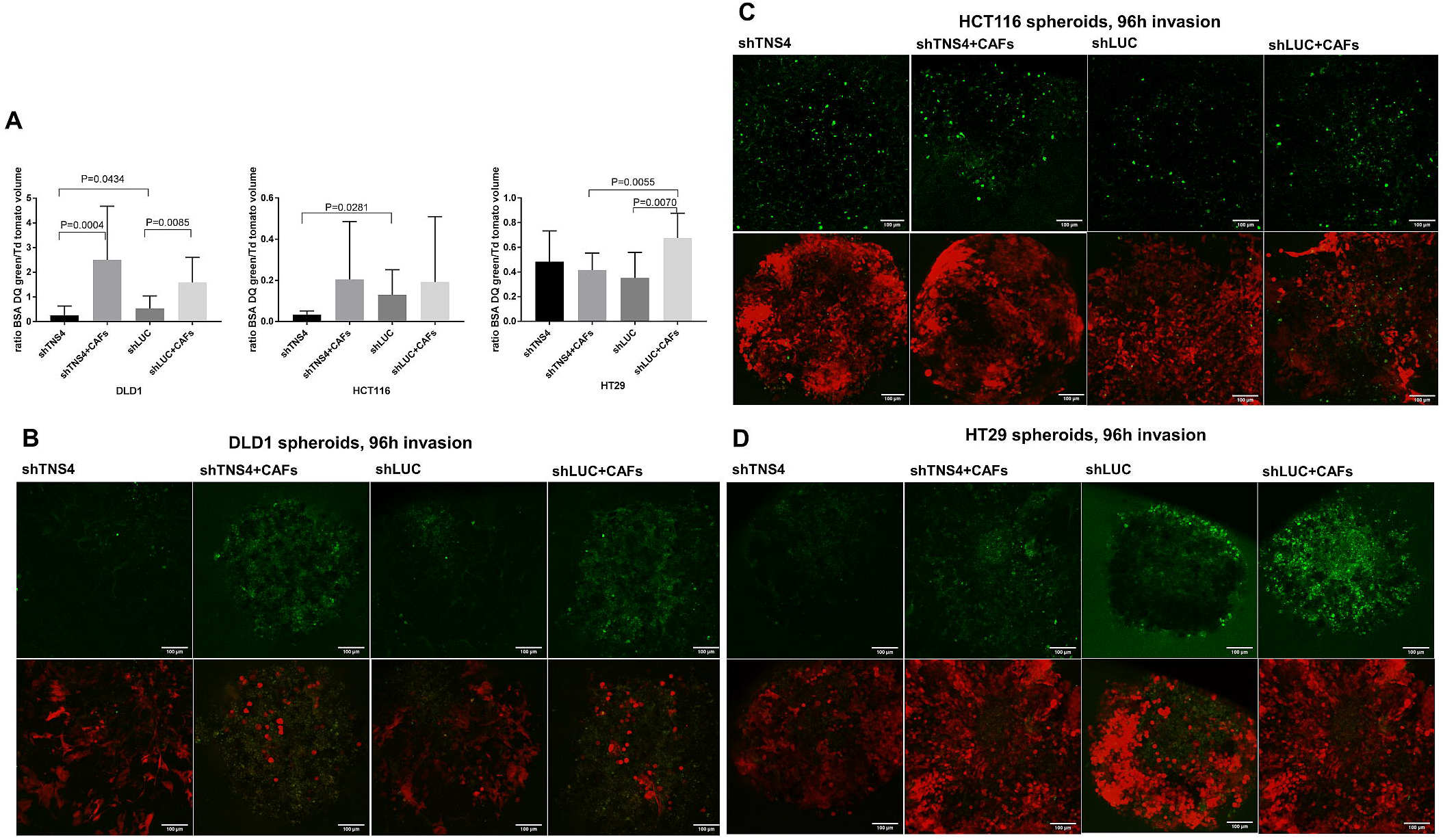
A) 96h measurement of extracellular matrix degradation in 3D spheroid invasion assay using DQ BSA green. Mann-Whitney t-test was used for comparisons between 2 groups of non-parametric data. Exact P-values shown are representative of 2 independent experiments. Representative images of Z-stack slice of 3D spheroids for DLD1 (B), HCT116 (C) and HT29 (D). Upper row depicts DQ BSA green fluorescence and lower row a projection of the maximum intensity of all slices in the confocal microscopy Z-stack.

### TNS4 knockdown sensitizes CRC cells to EGFR inhibition treatment

We and others have shown that TNS4 is an effector of the EFGR-KRAS-MAPK signalling pathway. This prompted us to investigate whether removing TNS4 could affect the activity of anti-EGFR drugs such as Gefitinib (which inhibits the kinase domain of EGFR). We investigated this in both 2D and 3D models.

Overall, we observed that knockdown of TNS4 resulted in increased chemosensitivity to gefitinib in all cell lines. This was seen in both assays, but the effect was more pronounced in 3D spheroids (Figure 6A and B). Gefitinib treatment alone in shTNS4 transduced cells reduced expression of EGFR, RAS and TNS4, thereby confirming that TNS4 forms part of this signalling pathway (Figure 6C and Supplementary Figure 2). Interestingly, TNS4 knockdown resulted in higher EGFR levels than in shLUC controls suggesting a negative feedback loop between TNS4 and EGFR),

**Figure 6.**
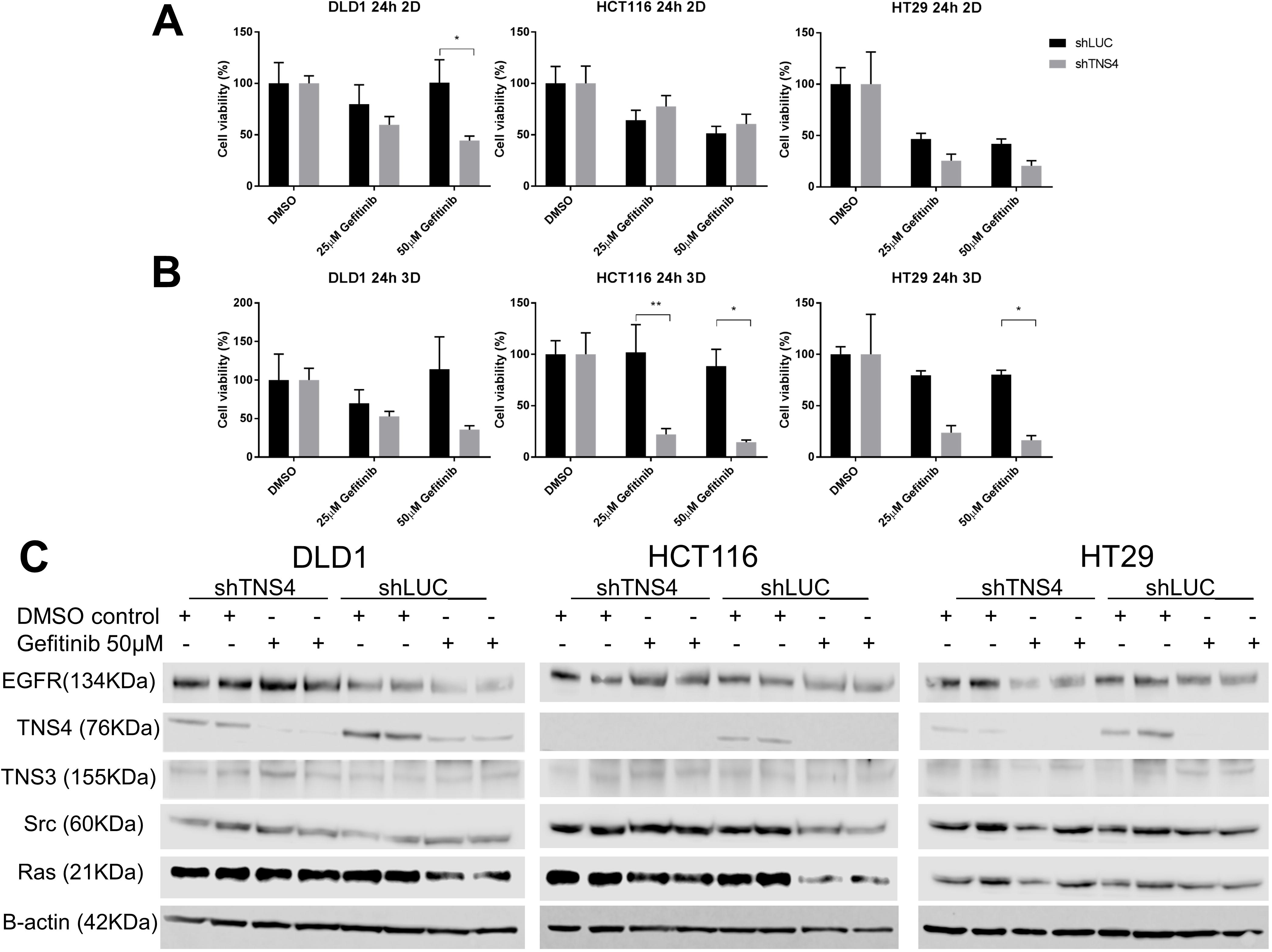
Chemosensitivity of shTNS4 and shLUC CRC cell lines in 2 D (A) and 3 D (B) conditions. Western blot protein detection of EGFR, TNS4, TNS3, Src and Ras in shTNS4 and shLUC CRC cells, treated with Gefitinib 50µM for 24h and respective controls, in two samples obtained from independent experiments. P-values indicating statistically significant differences are marked by *(P<0.05) or **(P<0.01).

## Discussion

Tns4 overexpression has been reported in a number of cancers [6, 8, 24, 25] although in colorectal cancer *in vitro* studies have not shown a role for Tns4 in cell proliferation. These were always performed in 2D settings, omitting the presence of an enriched tumour microenvironment with increased cell-to-cell interactions and supporting stromal cells. Only recently it has been reported that in HeLa cells, TNS4 transient silencing reduced cell growth, even though its overexpression had not impact on cancer cell proliferation [26]. However, analysis of 3D models does take longer and thus we used lentiviral transduction with shRNA oligos to produce stable TNS4 knockdowns. This allowed spheroid growth to be monitored for 7 days.

We firstly tested for changes in adhesiveness to both acellular stroma composed of collagen type I and CAFs followingTNS4 knockdown. We observed that shTNS4 transfected cells show significantly reduced adhesion to CAFs and slightly decreased adhesion to collagen type I (Figure 2), This effect could be due to a disturbance in molecular mechanosensors where TNS4 is located [27]. Adhesion of CRC cells to CAFs might be mediated by the mutli-molecular adhesome of integrin B1 heterodimers. The cytoplasmic tails of integrins are bound to the C-terminus PTB domain of TNS4 which in turn forms a complex with several other proteins including actin. [28–31]. Therefore, it is possible that removing TNS4 protein by gene knockdown could disturb the stability of the integrin B1 adhesome and its ability to bind to CAF-secreted fibronectin.

We next tested whether TNS4 expression could affect cell proliferation in spheroid models. Interestingly it appears to vary according to the spheroid morphology produced. In SW620, SW480 and LS1034, which naturally produce irregular shaped and loose spheroids when grown in 3D, a reduction in TNS4 expression resulted in decreased cell proliferation. In contrast, HCT116 and HT29, which naturally produce tight spheroids, shTNS4 transduction caused a significant increase in proliferation (Figure 3). The data were consistent for both fluorescence measurements and spheroid volumetric analysis indicating that the observations were not technical artefacts. This change in cell proliferation observed in compact spheroids, could be related to the effects on spheroid compactness and cell adhesiveness by disturbing molecular mechanosensors at the focal adhesion complex where TNS4 is situated and altering FAK-related contact inhibition. An association between phosphorylated FAK and nuclear TNS4 expression has been previously reported in metastases of colorectal cancer cases [32], and it was suggested the two pathways may interact. It is possible that the fine balance existing between the extracellular matrix and cell-to-cell interactions, FAK activation could be influenced by a downregulation of TNS4 to alter cell proliferation [33].

When CAFs were added to the spheroid cultures, as documented in other studies, this supported growth of CRC cells thereby validating our 3D spheroid co-culture model [15]. Interestingly, in this context, a growth advantage is also seen for shTNS4+CAFs in cell lines forming compact spheroids (DLD1, HCT116 and HT29), but not in loose spheroids formed by SW620, SW480 and LS1034. This data could suggest a novel role for TNS4 in intercellular interactions. Integrins have been reported to form heterotypic complexes with adhesion molecules in adjacent cells and this may form part of a contact inhibition mechanism [28]. Alternatively, the explanation may lie in differences in the diffusion rates of CAF-secreted growth factors, such as TGFß [34], in tight versus loose spheroids.

When investigating invasion and matrix degradation of 3D spheroids, we observed that, in two cell lines, knockdown of TNS4 replicated the data seen in 2D analysis (Figure 5). Validating the effects of TNS4 in these more sophisticated models supports the role of TNS4 in carcinogenesis. When CAFs were added to the 3D spheroids, as expected, increased invasiveness was observed. This effect has been explained by CAFs-directed assembly of fibronectin and induction of integrins αV and β3. [35]. Furthermore, CAFs have also been described to increase contractility and traction forces in the ECM, mediated by non-muscle myosin II and PDGFα, which are then transmitted through α5β1 integrin to produce an alignment of the fibronectin matrix. Aligned fibronectin matrix features in invasion sites in prostatic and pancreatic carcinomas [36]. The effect of TNS4 knockdown in the co-culture model was broadly in-line with the effect seen in the 3D monoculture in as much as there was a reduction in matrix degradation but it wasn’t quite as marked. Interestingly, in HT29, whilst the monoculture had shown no effect following TNS4 knockdown, in the 3D co-culture, invasion was significantly reduced in shTNS4+CAFs spheroids when compared to shLUC+CAFs (Figure 5A and D). These observations were made on several repeated experiments although we do not currently have an explanation, but suspect it could be due to the wild-type status of KRAS gene in HT29 and its different EGFR signalling potency.

Given our previous data demonstrating the involvement of TNS4 in EGFR/KRAS/MAPK signalling [4, 37, 38], we have used our model to test this in 3D spheroids. We also tested whether TNS4 could modulate the activity of anti-EGFR therapies by using gefitinib, which is an inhibitor of the EGFR kinase domain. Our results show a reduction of TNS4 levels with gefitinib treatment (Figure 6C), thereby confirming TNS4 is part of this pathway and suggesting the blockade of EGFR tyrosine kinase domain could lead to TNS4 increased degradation.

It has been shown that TNS4 stabilizes EGFR by reducing its ligand-induced degradation via a reduction in its ubiquitination, but not EGF-induced tyrosine phosphorylation of EGFR [39].. In shTNS4 spheroids, EGFR would be more unstable and easily degraded upon ligand-binding. Lower levels of active EGFR would also lead to increased sensitivity to its inhibitor gefitinib, which we have observed in DLD1 and HT29 (Figure 6A,C and Supplementary Figure 2). By performing gefitinib chemosensitivity test using shTNS4 and shLUC cells, we have observed a reduction in cell viability with TNS4 knockdown in both 2D and 3D conditions (Figure 6A and B). This supports the notion that TNS4 is an important downstream component of the EGFR signalling pathway and raises the possibility of an anti-TNS4 therapy being used in combination with anti-EGFR therapies. A recent study has also demonstrated that TNS4 is significantly downregulated by cetuximab (anti-EGFR immunotherapy) and put forward the possibility that TNS4 suppression could be used in cetuximab-refractory CRC patients with *KRAS* activating mutations [40].

In summary, we have hereby used a novel methodology to mimic an enriched 3D colorectal tumour microenvironment and demonstrate a role for TNS4 in the dynamics of cell adhesion, 3D spheroid proliferation, invasion and EGFR inhibitor resistance. These results suggest TNS4 has a valuable role as an effector in early stage metastasis, likely associated with its mechanosensing properties and involvement in heterotypic interactions at the focal adhesion complex. Furthermore, TNS4 inhibition might improve the efficacy of anti-EGFR therapy, by increasing its chemossensitivity.

## Acknowledgments

The Authors acknowledge financial support from the Nottingham Molecular Pathology Node – Medical Research Council, United Kingdom grant RA4852.

## Data availability statement

The data that support the findings of this study are available from the corresponding author upon reasonable request.

## Author Contributions

TPR conducted all the experimental work, analysed data and drafted the manuscript. SS established the CAFs line and optimized the spheroid co-culture system, MI reviewed and edited the manuscript. All authors read and approved the final manuscript.

---

**Supplementary Table 1:**
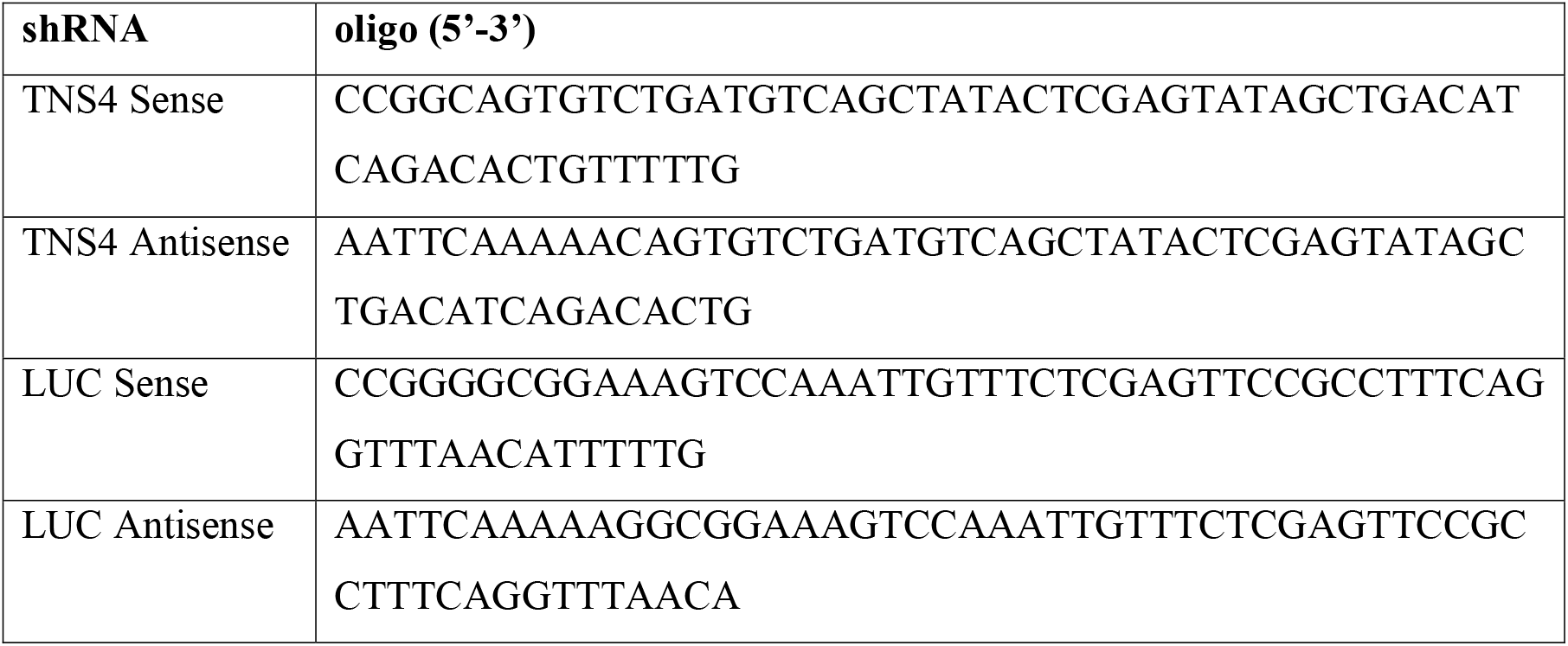
shRNA oligonucleotides for TNS4 and LUC.

**Supplementary Table 2.**
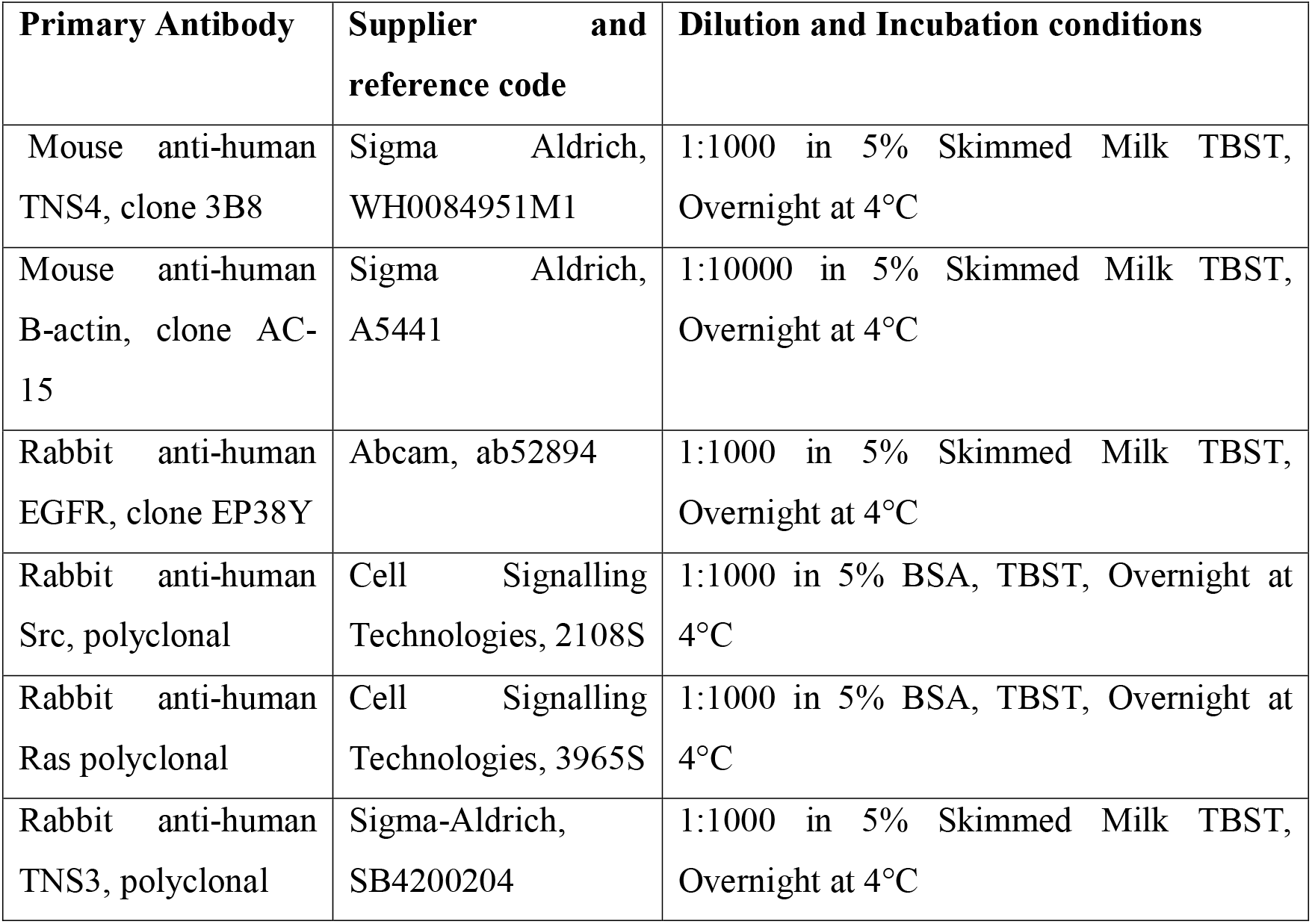
Primary antibodies, suppliers, dilution and incubation conditions used for western blot.

| **Primary Antibody** | **Supplier and reference code** | **Dilution and Incubation conditions** |
| --- | --- | --- |
| Mouse anti-human TNS4, clone 3B8 | Sigma Aldrich, WH0084951M1 | 1:1000 in 5% Skimmed Milk TBST, Overnight at 4°C |
| Mouse anti-human B-actin, clone AC-15 | Sigma Aldrich, A5441 | 1:10000 in 5% Skimmed Milk TBST, Overnight at 4°C |
| Rabbit anti-human EGFR, clone EP38Y | Abcam, ab52894 | 1:1000 in 5% Skimmed Milk TBST, Overnight at 4°C |
| Rabbit anti-human Src, polyclonal | Cell Signalling Technologies, 2108S | 1:1000 in 5% BSA, TBST, Overnight at 4°C |
| Rabbit anti-human Ras polyclonal | Cell Signalling Technologies, 3965S | 1:1000 in 5% BSA, TBST, Overnight at 4°C |
| Rabbit anti-human TNS3, polyclonal | Sigma-Aldrich, SB4200204 | 1:1000 in 5% Skimmed Milk TBST, Overnight at 4°C |

Supplementary Figure 1 – Morphology of spheroids from different colorectal cancer cell lines. Compact and round spheroids are produced by cell lines DLD1, HCT116 and HT29 and volume estimations can be taken. Irregular, loose spheroids or aggregated cells are produced by LS1034, SW480 and SW620 and their growth rate is best captured by measurement of fluorescence in stably labelled cells, for example by lentiviral transduction of fluorescent protein encoding viral vectors.

Supplementary figure 2- Densitometry for WB shown in figure 6, showing relative ratios of EGFR, Cten, TNS3, Src and Ras protein levels to the loading control B-actin in the following CRC cell lines: DLD1 (A), HT29 (B) and HCT116 (C). A statistical analysis of the differences observed in the densitometry values was not carried out as only 2 independent biological samples were used for the western blots.

