## Supplementary figures and images for "Investigating TNS4 in the Colorectal Tumour Microenvironment Using 3D Spheroid Models of Invasion"

### Supplementary Figure 1

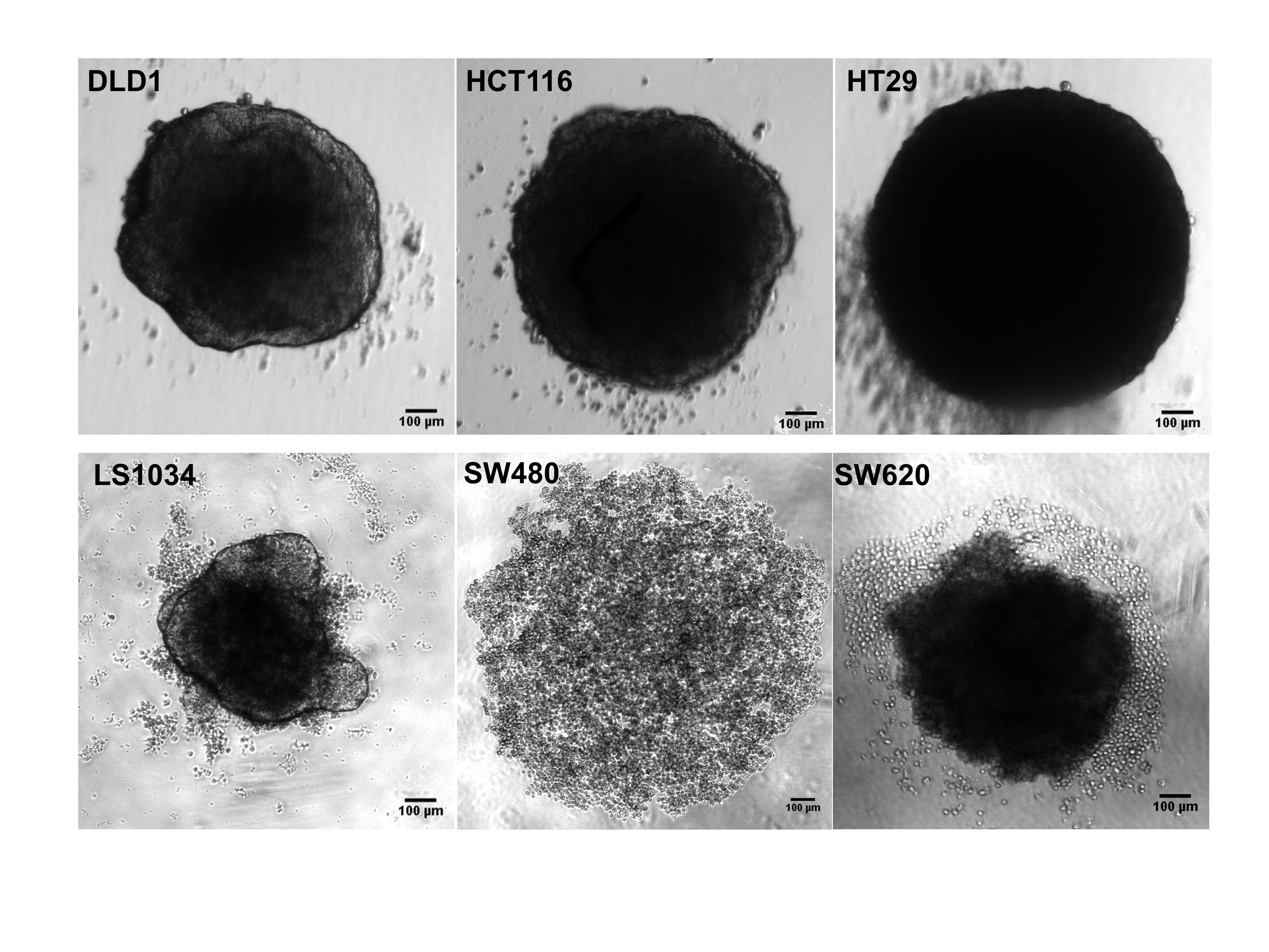

### Supplementary Figure 1

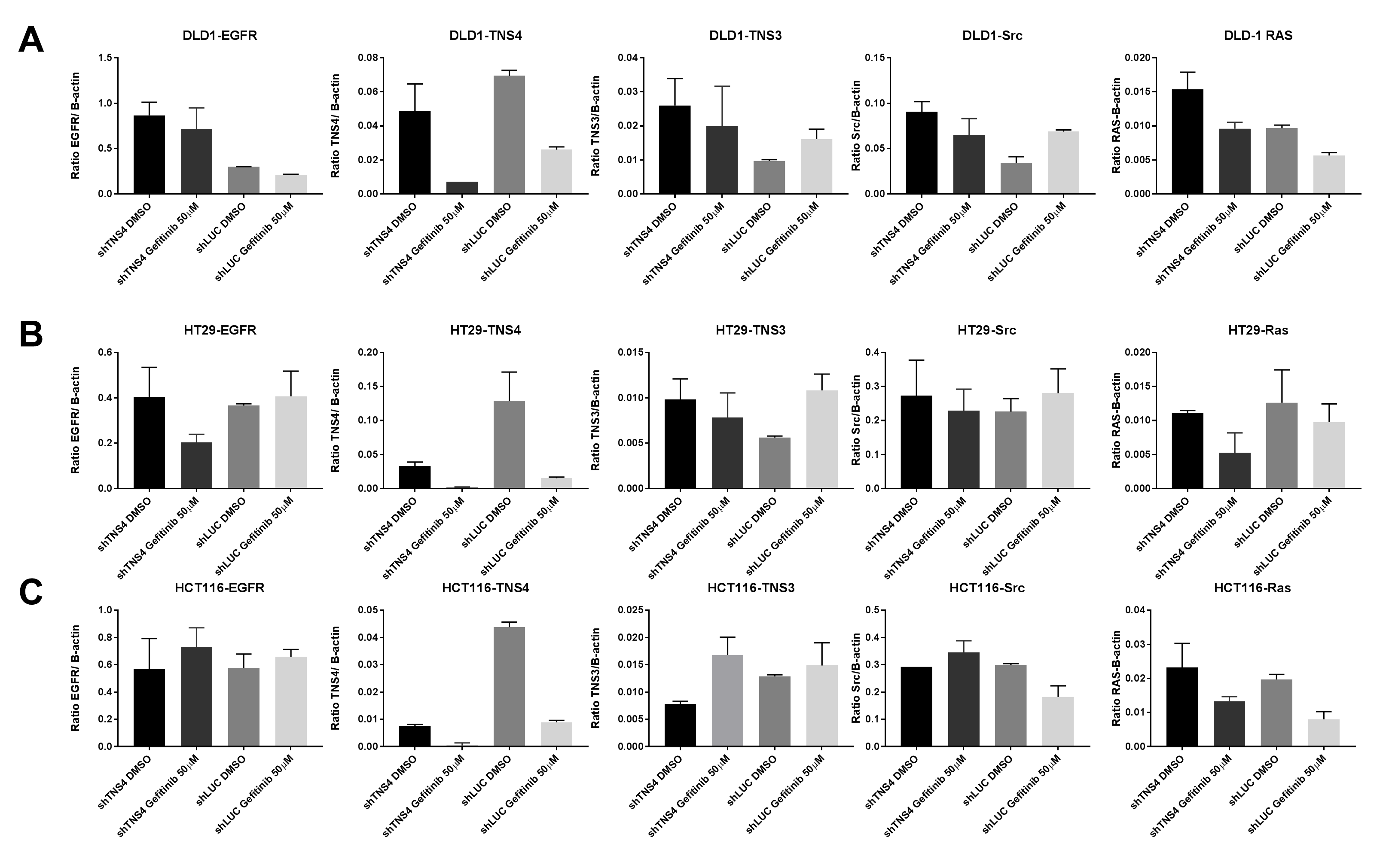
